# Structural Ensembles of Ribonucleic Acids From Solvent Accessibility Data: Application to the S-Adenosylmethionine (SAM)-Responsive Riboswitch

**DOI:** 10.1101/2020.05.21.108498

**Authors:** Jingru Xie, Aaron T. Frank

## Abstract

Riboswitches are regulatory ribonucleic acid (RNA) elements that act as ligand-dependent conformational switches. In the apo form, the aptamer domain, the region of a riboswitch that binds to its cognate ligand, is dynamic, thus requiring an ensemble-representation of its structure. Analysis of such ensembles can provide molecular insights into the sensing mechanism and capabilities of riboswitches. Here, as a proof-of-concept, we constructed a pair of atomistic ensembles of the well-studied S-adenosylmethionine (SAM)-responsive riboswitch in the absence (-SAM) and presence (+SAM) of SAM. To achieve this, we first generated a large conformational pool and then reweighted conformers in the pool using solvent accessible surface area (SASA) data derived from recently reported light-activated structural examination of RNA (LASER) reactivities, measured in the −SAM and +SAM states of the riboswitch. The differences in the resulting −SAM and +SAM ensembles are consistent with a SAM-dependent reshaping of the free landscape of the riboswitch. Interestingly, within the −SAM ensemble, we identified a conformer that harbors a hidden binding pocket, which was discovered using ensemble docking. The method we have applied to the SAM riboswitch is general, and could, therefore, be used to construct atomistic ensembles for other riboswitches, and more broadly, other classes of structured RNAs.

## INTRODUCTION

Changes in conformational equilibria – in response to changes in physiochemical conditions within the cell – underlie the biological function of many ribonucleic acids (RNAs). Such changes may include changes in temperature, pH, or the absence/presence of binding partner(s). This ability of RNA to respond to changes in local cellular conditions is best exemplified by riboswitches, which are cis-acting regulatory RNA elements located in the 5’-untranslated (UTR) region of mRNAs that change their conformation upon binding to specific ligands.(1–3) This conformational change either sequesters or releases sequence-motifs that, in turn, activate or deactivate transcription or translation. The aptamer domains of riboswitches bind to their cognate ligands with high specificity and confer the RNA with its sensing capabilities. As such, understanding the structure of the aptamer domain is critical to understanding relationships between the conformational equilibria of aptamers and the sensing capabilities of riboswitches. Mounting evidence suggests that the aptamer domain of riboswitches exhibits varying degrees of structural plasticity. Therefore, characterizing the structural ensemble, comprised of the set of conformers that are accessible to a riboswitch under a specific set of conditions, is a critical step in describing and then rationalizing its response to cognate and non-cognate ligands. Analysis of such ensembles can reveal the existence of alternate binding pockets that may facilitate binding of non-cognate ligands, as well as novel allosteric sites that may facilitate ligand binding away from the site bound by cognate ligands.

Solution techniques can be used to probe the equilibrium conformations of RNAs by providing access to structure-dependent, ensemble-averaged measurements that can, in principle, be used to infer structural ensembles that capture the conformational distribution of an RNA under a specific set of conditions. For instance, chemical probing experiments can be used to identify reactive sites in RNA, both *in vitro* and *in vivo* (4). Because the sites that are most “reactive” tend to be solvent-exposed, the reactivities obtained from these experiments provide an indirect “read-out” of the local solvent accessibility across the ensemble of structures populated by the RNA. The ensemble-averaged reactivities derived from light-activated structural examination of RNA (LASER) experiments, in particular, have been shown to correlate strongly with solvent accessible surface area (SASA) of the C8 atom in purine residues (5), suggesting that they might be useful for constructing such ensembles. Using SASA data derived from highly accessible measurements is advantageous as it provides a fast and efficient route to access structural ensembles that can be used to infer functionally relevant environmental responses in RNA, and to yield individual conformers for the structure-guided design of therapeutics.

To infer structural ensembles from ensemble-averaged experimental data like SASA, one of two strategies can be employed: restraining or reweighting.(6) Restraining involves carrying out molecular dynamics simulations with a force field augmented with restraint terms that ensure that the simulated ensemble-averaged observables matches the experimental measurements (7–9). Reweighting, on the other hand, is a post-processing approach that involves assigning weights to conformers within an ensemble, so that the ensemble-averaged observables computed using these weights match the measured observables (10–12). As a post-processing approach, reweighting has the advantage of being computationally efficient and could be used to generate multiple ensembles from multiple sets of experiment data (possibly measured under differing conditions) from the same set of structures without requiring additional simulations.

Here, employing ensemble reweighting, we used C8-SASA derived from LASER experiments for S-adenosylmethionine (SAM)-responsive riboswitch in the absence and presence of SAM, to infer atomistic ensembles of the aptamer domain of the SAM riboswitch. The ensembles recapitulated global and local features that were consistent with previous experiments and simulations. Within the ligand-free ensemble, we identified conformers of the SAM riboswitch that possess alternate binding pockets that are predicted to accommodate small-molecule ligands besides the cognate SAM ligand. We envision that the approach we used for the aptamer domain of SAM riboswitch could also be used to model the ensemble of other structured RNAs.

## MATERIALS AND METHODS

### Benchmarking the ability of SASA to recover ensembles

#### Benchmark data set

To benchmark the inherent ability of SASA data to recover atomistic ensembles of RNA, we first compiled a data set of 45 RNA molecules, with lengths ranging from 14 to 53 nucleotides (Table S1). For each RNA, its atomic structure (crystal or NMR structure) was downloaded from Protein Data Bank(13), which was then used to generate a set of diverse conformations using FARFAR (Fragment Assembly of RNA with Full Atom Refinement) (14, 15) and KGSrna (Kino-geometric sampling for RNA) sampling(16). 1000 conformations were generated using each method, resulting in a simulated conformational pool of 2000 conformations. For each RNA, we sampled 200 structures from this conformational pool to form a **decoy ensemble** that is uniformly distributed in RMSD space. The **target ensembles** were then formed by selecting the native structure and a set of structures from the decoy ensemble within an RMSD cutoff to the native structure. The RMSD cutoff, or ensemble width, was varied between 2.0 to 11.0 Å in increments of 1.0 Å to assess the reweighting power of SASA for various target ensembles. From each *target ensemble*, ensemble-averaged SASA were calculated for each C8 atom in the purine residues and used as *target data* to reweight the *decoy ensemble* using the Bayesian maximum entropy (see below).(17, 18)

#### Bayesian maximum entropy (BME) method

We cast the problem of constructing ensembles using SASA data as a reweighting problem, in which weights are applied to conformers in the decoy ensemble conditioned on the ensemble-averaged SASA data associated with the target ensemble. Formally, we sought to reweight the decoy ensemble using a set of optimal weights {*w*_*i*_}, with *w*_*i*_ being the weight for a conformer *i* in the decoy ensemble, such that ensemble-averaged SASA from the reweighted ensemble best matches the target SASA. Within the BME scheme (17, 18), the problem of finding the optimal {*w*_*i*_} is formulated as a constrained optimization problem with the entropy of weights being the objective function. Specifically, we attempted to find the *{w*_*i*_*}* that

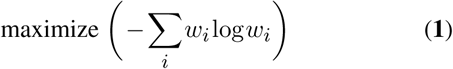

subject to

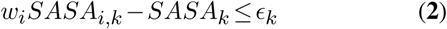

where *SASA*_*i,k*_ is the SASA of atom *k* in conformer *i, SASA*_*k*_ is the ensemble-averaged SASA of atom *k* in the target ensemble, and *ϵ*_*k*_ is the error tolerance sampled from a Gaussian distribution (*p*(*ϵ*_*k*_)):

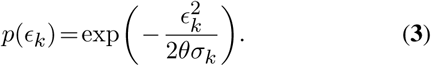

where *σ*_*k*_ is the standard deviation calculated from the target SASA, and *θ* is a factor that scales *σ*_*k*_. For each RNA in our benchmark data set, we utilized the BME approach to reweight the decoy ensemble using C8-SASA data from each of the 10 distinct target ensembles, which differ in terms of the RMSD cutoff to the native structure (see above). Furthermore, for each decoy and target ensemble pair, we ran 500 trials with random initialization of *ϵ*_*k*_. The reweighted weights were averaged over all trials to get stable and reliable results, which minimizes the effect of randomness introduced by the sampling of *ϵ*_*k*_. Therefore, in total, we carried out 225000 (45 10 500) reweighting experiments; in each case, we used *θ* = 1.0 (Eq. **3**).

#### Comparing Ensembles

To assess the performance of the C8-SASA BME reweighting on the benchmark data set, we compared the reweighted and decoy ensembles to the target ensembles. Firstly, the atomic density maps for decoy, target, and reweighted ensembles were calculated using GROma*ρ*s(19), a GROMACS-based density map analysis tool. Secondly, to visually compare the difference, we computed the difference density maps between target and reweighted ensembles and between target and decoy ensembles. The difference maps were then rendered as volume maps using PyMOL (version 2.3.4). Thirdly, we calculated the cross-correlation between the decoy and target density maps (*CC*_*DT*_), and the cross-correlation between the reweighted and target density map (*CC*_*RT*_). The cross-correlation is the global correlation as implemented in GROma*ρ*s(19). Finally, we calculated, which we defined as the ratio between two cross-correlation values, that is,

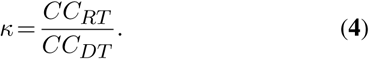

Values of κ > 1 correspond to instances in which the atomic density maps of the reweighted ensemble more closely resembled (i.e., were more highly correlated with) the target ensemble than the decoy ensemble (i.e., *CC*_*RT*_ *>CC*_*DT*_).

### Constructing ensemble of the SAM riboswitch

To construct structural ensembles of the SAM riboswitch in the absence (-SAM) and presence (+SAM) of SAM, we first generated a conformational pool using the conformational sampling tool, KGSrna. Using KGSrna, we generated a total of 32,000 conformers starting from the crystal structure of −SAM (PDBID: 3IQP(20)) and +SAM (PDBID: 2GIS(21), 3IQR(20)) states. Next, from each conformer in the conformational pool, C8-SASA of all purine residues were computed using FreeSASA (22). Using LASER-derived C8-SASA as the target data, we then reweighted the conformational pool. To estimate C8-SASA from LASER reactivity data, we fit LASER reactivity data(5) to FreeSASA-computed C8-SASAs for +SAM crystal structure (PDBID: 2GIS) to obtain a linear function that maps LASER reactivity to C8-SASA (21). The fit was then used to estimate C8-SASA for the −SAM and +SAM states from available −SAM and +SAM LASER reactivity data, respectively (5). LASER-estimated C8-SASA was then used as the mean of target distribution to reweight the 32,000-membered conformational pool using the BME reweighting (see above).

### Cavity mapping and docking experiments

For each of the four highest weighted conformers in the −SAM ensemble, we carried out cavity mapping to identify sites on the surface that might facilitate interactions with small-molecule ligands. To achieve this, we utilized the two-sphere cavity mapping method implemented in the rbcavity program within the rDock modeling suite (23). For cavity mapping, we set the maximum number of cavities to 10 and the minimum cavity volume to 50 Å^2^. The resulting cavities were visualized in PyMOL (version 2.3.4). Next, we carried out a small-scale *in silico* screening by docking 500 small, drug-like molecules into the cavities identified using rbcavity. These 500 small molecules corresponded to a small library of drug-like compounds obtained from the ZINC library (24). To identify the most conformationally selective compounds in the library, we first computed the selectivity index, *γ*_*i,j*_, which we defined as

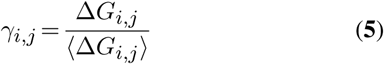

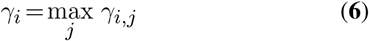

Here *i* runs over the compounds in the library, *j* runs over the conformers (docking receptors), Δ*G*_*i,j*_ is the docking score for compound *i* docked onto conformer *j*, and ⟨Δ*G*_*i,j*_⟩ is the average docking score for compound *i* across all *j* conformers. For each compound, we assign a selectivity (*γ*_*i*_) as the maximum of the set of selectivity indices {*Δ*_*i,j*_}. As defined, compounds with high *γ*_*i*_ correspond to those that have a docking score (Δ*G*_*i,j*_) on a given conformer that is significantly more favorable (negative) than the average docking score (⟨Δ*G*_*i,j*_⟩); such a compound was identified as being a “conformationally selective” compound.

## RESULTS

### Benchmark results suggest that SASA-reweighted ensembles more closely resemble targets ensembles than decoy ensembles

In this study, we set out to infer solution structural ensembles of the *Thermoanaerobacter tengcongensis* SAM riboswitch in the absence (-SAM) and presence (+SAM) of SAM, using available LASER reactivity data(5). The SAM-responsive riboswitch, which regulates genes that control the metabolism of sulfur and methionine, consists of four helices (P1, P2, P3, and P4) that are connected by a four-way junction, with the SAM binding site located between P1 and P3 (25). Previous studies support the hypothesis that the −SAM state of the riboswitch samples a wide range of conformational states in solution, including bound-like conformers (20).

Before constructing these SAM ensembles, we benchmarked the reweighting power of C8-SASA data. In particular, we explored the sensitivity of the SASA reweighting scheme to both error in the SASA data and the width of the target ensemble, from which we computed the ensemble-average SASA data used for reweighting. To carry out this benchmarking, we first generated pairs of decoy and simulated target ensembles. Then, using ensemble-averaged C8-SASA from the target ensemble, we reweighted the decoy ensemble. Visual inspection of the difference atomic density maps of the reweighted ensembles relative to their target ensembles revealed that, in general, the C8-SASA reweighted ensembles exhibited smaller residual densities than the initial decoy ensembles (Figure 1a, b, S4). This observation suggests that the reweighted ensembles tend to more closely resemble the target than did the decoy ensembles. Shown in Figure 1c are plots of *κ* (Equation **4**), the ratio of the cross-correlation between the atomic density maps of the reweighted ensemble (*CC*_*RT*_) and the decoy ensemble (*CC*_*DT*_) relative to the target ensemble. Results are shown as a function of noise-level in the target SASA and the width of the target ensembles. As defined, when the correlation between the reweighted and target ensemble is higher than that of the decoy and target ensembles, *κ* > 1. As might be expected, the lower the level of noise added to the target data and the narrower the width of the target ensemble, the higher the values of *κ*. Accordingly, *κ* is > 1 when the width of the target ensemble is *<*= 4.2 Å and the noise-level is *<*= 1.3 Å^2^, which suggests that under such conditions, the C8-SASA data can be used to bring the decoy ensembles into better correspondence to the target ensembles.

**Figure 1.**
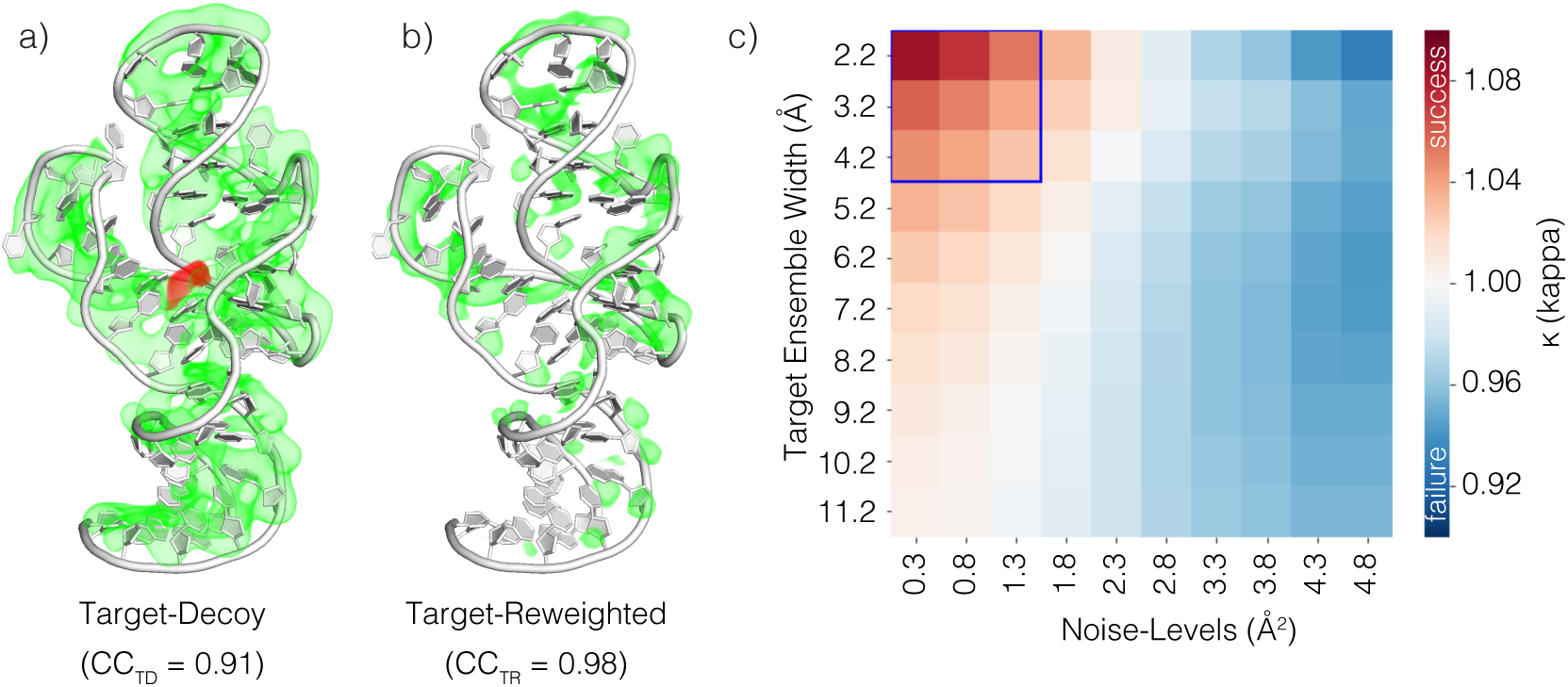
Benchmarking results. Shown are comparisons between the difference-atomic maps between (a) the target and decoy ensembles and (b) the target and reweighted ensembles for the CR4/5 domain of medaka telomerase RNA (PDBID: 2MHI), one of the 45 RNAs in our benchmarking data set (Supporting Information). (c) Heatmap of *κ* as a function of the width of the target ensembles and the noise-level in the corresponding target C8-SASA data. Ensemble width is defined as the maximum RMSD between a structure in the target ensemble and the native structure. The noise-level in target C8-SASA is simulated by adding random noise to the target ensemble-averaged C8-SASA. The absolute value of the noise was sampled from an exponential distribution with noise level as the scale parameter (or average noise). The map shown here is the average of *κ* values over all the RNAs in our data set. Note that for some RNA ensembles, the BME algorithm failed to converge. Accordingly, the averaging is performed on successful reweighting only.

### SASA-based ensembles of the SAM riboswitch are consistent with reshaping of the conformational pool in the presence of SAM

Next, using SASA data derived from LASER experiments, we constructed ensembles for the −SAM and +SAM states of the aptamer domain of the SAM riboswitch. To generate the ensembles for the −SAM and +SAM states of the riboswitch, we generated a diverse pool of conformers and computed the C8-SASA for purine residues, which correspond to the site in RNA probed by LASER experiments. We then reweighted the pool of structures using the Bayesian maximum entropy (BME) reweighting method,(18) using the C8-SASA predicted from LASER reactivity measured in - SAM and +SAM states as the target data, respectively(5). Briefly, to generate the target data, we converted LASER reactivities to C8-SASA by fitting LASER reactivities of the +SAM to C8-SASA computed for the crystal structure of the +SAM. The mean error in the fit was 1.27 Å^2^ (Supporting Information). Based on the benchmarking results presented above (Figure 1c), we expect that ensembles reweighted using C8-SASA data with an inherent error 1.27 Å^2^ should be closer to the “true” ensembles than the initial conformational pool. Shown in Figure 2 is the comparison between the average structure in LASER/SASA-derived ensembles for the −SAM (Figure 2a) and +SAM (Figure 2b) states. The RMSD of the average ensemble structures were only 1.30 Å consistent with X-ray crystallography result that the −SAM and +SAM structures were almost identical (RMSD = 0.52 Å). Despite the similarity of the average −SAM and +SAM structures, we did observe some subtle differences in the - SAM and +SAM ensembles along the distributions of the distance between residue 47 and residue 90, which we used as the reaction coordinate to describe the openness of P1 relative to P3 (Figure 2c). Comparison of the −SAM and +SAM distributions revealed that the mode of distribution shifts from 16.8 to 13.0 Å when going from the −SAM to the +SAM state, consistent with the −SAM state having a preference, relative to the +SAM state, for the open P1-P3 state.

**Figure 2.**
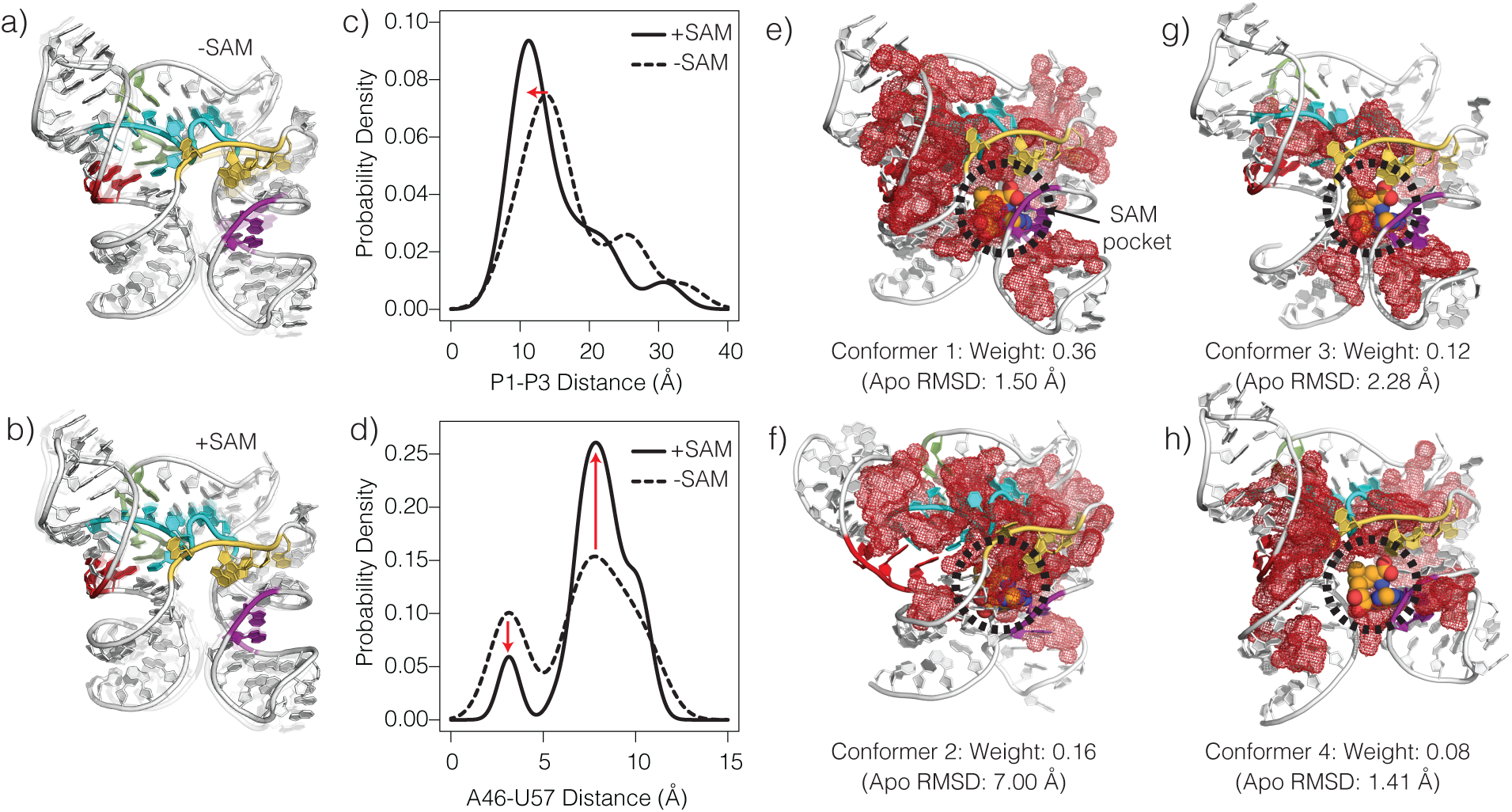
The LASER/SASA-derived SAM ensemble. Shown are the average structures in the LASER/SASA-derived ensemble of the −SAM (a) and +SAM (b) states of the SAM riboswitch. Shown in (c) is the distribution of the distance between P1 and P3 helices for both the −SAM and +SAM states. Similarly, shown in (d) are the distribution of the distances between residues A46 and U57, which are paired in the −SAM state (A46/U57 closed) and unpaired (A46/U57 open) in the +SAM state. (e-h) The four highest weighted conformers in the −SAM ensemble. For reference, the SAM is overlaid onto the images. The red mesh highlights the cavities identified in each conformer.

Despite the global structure of the −SAM and +SAM being almost identical, an inspection of the crystal structures reveal that in the −SAM, A46 and U57 are base-paired (closed), whereas in the +SAM state, they are not base-paired (open); U57 instead forms a hydrogen bond with SAM in the +SAM state. To test whether our ensembles captured this subtle difference, we computed the distribution of the distance between A46-U57, which we used as a reaction coordinate to describe the extent to which A46-U57 were base-paired (Figure 2d). In contrast to the distribution of the P1-P3 distance (Figure 2c), we observed a more dramatic difference between the −SAM and +SAM (Figure 2d). The distribution of the A46-U57 distance in the LASER/SASA ensembles are consistent with −SAM sampling both the closed and open A46/U57 states, whereas +SAM predominantly existing in the open A46/U57 state (Figure 2d). The LASER/SASA-derived ensembles, therefore, support a mechanism in which SAM shifts the population of states toward the closed A46/U57 state (Figure 2d).

### The −SAM ensemble contains a conformer that is predicted to bind to small-molecules via a hidden pocket

Since their discovery, riboswitches have garnered interest as under-explored drug targets (26). Indeed, the recent discovery of the antibacterial small molecule, ribocil-B, which targets the flavin mononucleotide (FMN) riboswitch supports the notion that riboswitches are druggable RNA targets (27). As a result of this and related discoveries (28), the identification and design of small molecules that target riboswitches has become an active area of research. Because individual riboswitches have evolved to bind to a specific ligand, attempts to design compounds that target riboswitches have focused on identifying compounds that are analogs of the cognate ligand or compounds that can recapitulate its interaction pattern. Alternatively, one could envision identifying small molecules that bind to a riboswitch at a site other than that occupied by the cognate ligand. Once identified, these alternative sites can be targeted using structure-based methods like molecular docking (29).

To illustrate how one might attempt to detect such site computationally, we applied ensemble docking to the highest weighted conformers in our −SAM ensemble. First applied the two-sphere cavity mapping method to the four highest weighted conformers in the −SAM ensemble; these four conformers have a cumulative weight of >0.70 (Figure 2e-h). Across the four conformers, we observed vast variation in both the location and size of the cavities (Figure 2e-2h). Next, we next docked a set of 500 small, drug-like molecules (chosen from the ZINC library(24)) onto each of the four conformers. Because we were particularly interested in identifying dockable binding pockets, into which small molecules can fit, we focused our analysis on the short-range van der Waals (VDW) contribution to the binding free energy. Shown in Figure 3a are the distributions of the non-polar contribution to the binding free energy (Δ*G*) estimated using the rDock function. The mean Δ*G* across all four conformers was −24.0 kcal/mol, indicating that in general the small molecules in the library were capable of forming favorable interaction with −SAM conformers 1-4. Overall, however, conformer 2 exhibited Δ*G* values than the other conformers (Figure 3a). Finally, of the screened molecules, we computed *γ*_*i*_, the selective index (Equation **6**), to identify those that exhibit conformational selectivity across the four conformers. Interestingly, the six most selective compounds in our library all exhibited a preference for conformer 2 (Table 1). Moreover, in this conformation, all six compounds are predicted to bind to the SAM riboswitch at the same binding site, which is located away from the binding pocket occupied by SAM in the +SAM crystal structure. Intriguingly, the pocket occupied by these molecules is near a set of conserved residues that participate in a base-triple that, though far away from the SAM binding site, have been shown to exhibit the strongest SAM-dependent stabilization (20). In conformer 2, however, the base-triple is absent (Figure 3b), which results in the formation of the binding cavity that these compounds occupy. The pocket that these molecules occupy, therefore, represents a “hidden” pocket that is absent from the −SAM and +SAM crystal structures, and which only emerged after conformational sampling. Within the pocket, the six selective compounds are predicted to form stacking interactions with a pair of conserved residues, A62, and C65 (Figure 3b; Figure S4). Collectively, these results suggest that the simulated −SAM ensemble contain conformers that harbor dockable pockets other than the pocket occupied by the cognate ligand, SAM.

**Table 1.** Docking scores of conformationally selective binders. For each, listed are the predictd binding free energy with conformer 1 (Δ*G*_1_), 2 (Δ*G*_2_), 3 (*G*_3_), and 4 (Δ*G*_4_). Also listed for each compound is *γ*_*i*_ *×*min({ΔG}), the product of selectivity index, and the lowest docking score across the four conformers. Here the the binding free energy correspond to the non-polar (Van der Waals) contribution estimated using the rDock scoring function.

| Compound | $\Delta G_1$ (kcal/mol) | $\Delta G_2$ (kcal/mol) | $\Delta G_3$ (kcal/mol) | $\Delta G_4$ (kcal/mol) | $\gamma_i \times \min(\{\Delta G\})$ (kcal/mol) |
| --- | --- | --- | --- | --- | --- |
| 1 | -18.5 | **-26.6** | -16.1 | -14.9 | -37.2 |
| 2 | -10.2 | **-24.0** | -15.3 | -13.5 | -36.4 |
| 3 | -17.0 | **-26.3** | -15.7 | -17.3 | -36.3 |
| 4 | -17.3 | **-26.1** | -16.6 | -15.6 | -35.9 |
| 5 | -22.8 | **-26.6** | -18.6 | -10.7 | -35.9 |
| 6 | -15.7 | **-27.5** | -20.2 | -21.7 | -35.4 |

**Figure 3.**
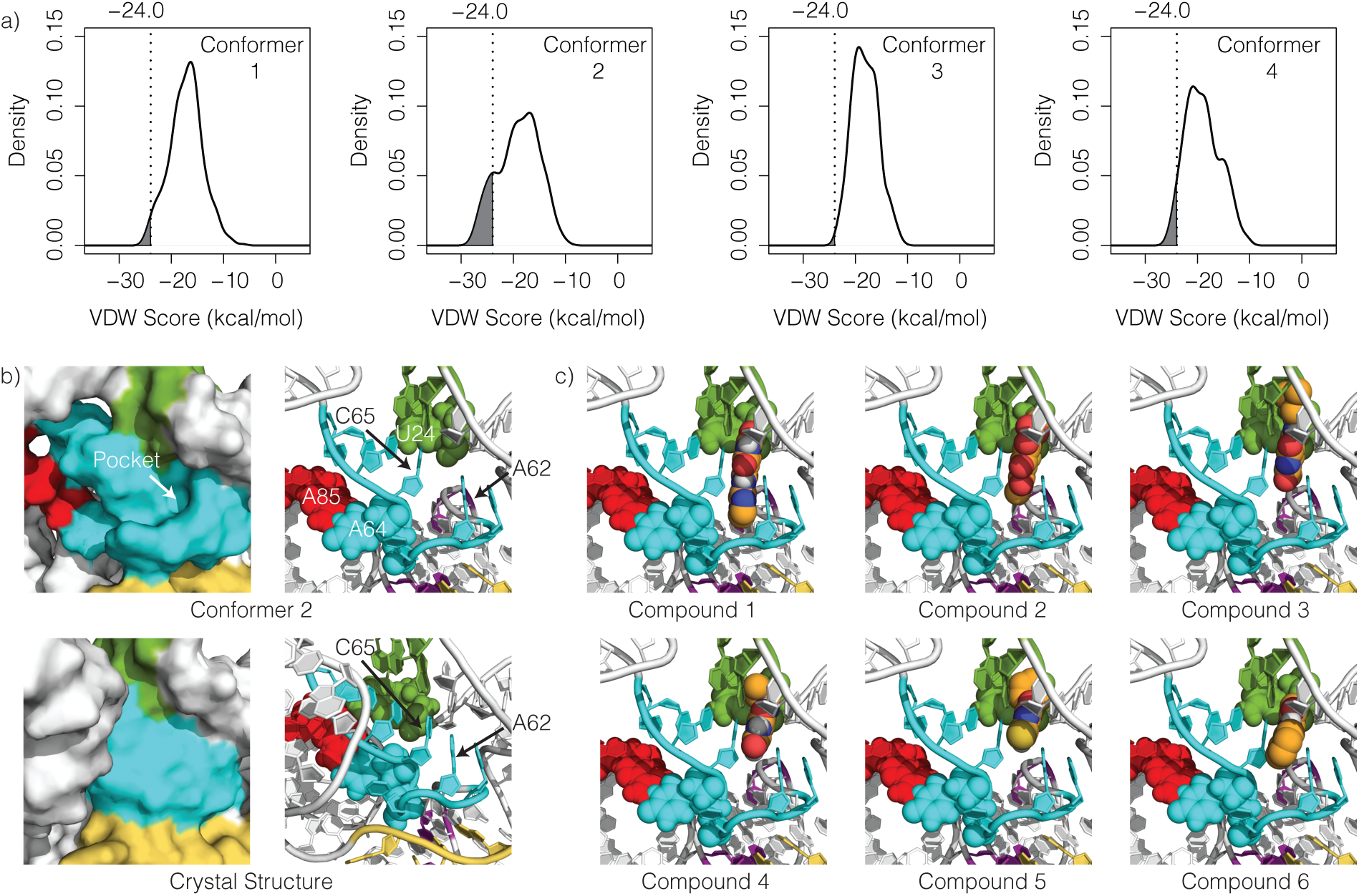
Ensemble docking. Distribution of docking scores across the conformer 1-4 in the −SAM ensemble. (b) A comparison between the binding site of the six most selective compounds in conformer 2 (top) and the corresponding site in +SAM crystal structure (bottom). The binding site is a hidden pocket, present in conformer 2 but absent in the +SAM crystal structure (bottom). Notably, the pocket features an increased nucleobases A62-C65 distance and the absence of the nearby U24-A64-A85 base-triple. (c) Poses of the six most selective small molecules docked onto conformer 2. All six compounds form stacking interactions with C65 and A62.

## DISCUSSION

Starting from crystal structures, we utilized conformational sampling and Bayesian maximum entropy (BME) reweighting to infer atomistic ensembles for the −SAM and +SAM states of the SAM riboswitch. The ensembles we generated were consistent with that −SAM state sampling a wider range of conformations relative to the +SAM state (Figure 2c,d), which was suggested by the existing biochemical and biophysical data (20). Interestingly, by docking a small library of compounds against the four highest weighted conformers in the −SAM ensemble, and then identifying selective binders in the library, we were able to locate what *appears to be* a “hidden” binding pocket in the −SAM riboswitch. Because the residues that line this pocket are highly conserved (Figure S4), it represents an ideal pocket for small molecule targeting. Future work will center around executing a more exhaustive search for compounds that target the hidden site we identified.

We note that the use of solvent accessible surface area (SASA) to infer the atomistic ensemble of the SAM-responsive riboswitch was predicated on the assumption that SASA inherently contains conformational restraining power. Recently, Madl and coworkers carried out solution NMR experiments in which they used solvent paramagnetic relaxation enhancements (sPRE) induced by the soluble, paramagnetic compound Gd(DTPA-BMA) to probe the structure of two benchmark RNAs.(30) Like the reactivities derived from chemical probing, the sPRE data provided an indirect “read-out” of local solvent accessibility across the equilibrium ensemble. They found that the inclusion of sPRE data during structural refinement significantly enhanced the quality of the resulting NMR ensemble.(30) Their findings, along with the results of our benchmarking studies, strongly suggest that experimentally-derived SASA contains sufficient restraining power to infer structural ensembles of RNAs. We envision, therefore, that the SASA-based reweighting approach we utilize in this study will emerge as a simple yet robust strategy for using experimentally-derived SASA data to infer atomistic ensembles. Such ensembles can then be used to generate or test structure-function hypotheses, as well as provide useful structural data to guide the discovery and design of RNA-targeting therapeutics.

## ACKNOWLEDGEMENTS

We thank Dr. Indrajit Deb for carrying the phylogenetic analysis for the class I SAM riboswitch and preparing the conservation map presented in the Supporting Information. We also thank Dr. Markos Koutmos, for many useful discussions about structural studies of riboswitches. The authors were funded by the University of Michigan research and computational startup funds.

---

**Figure S1:**
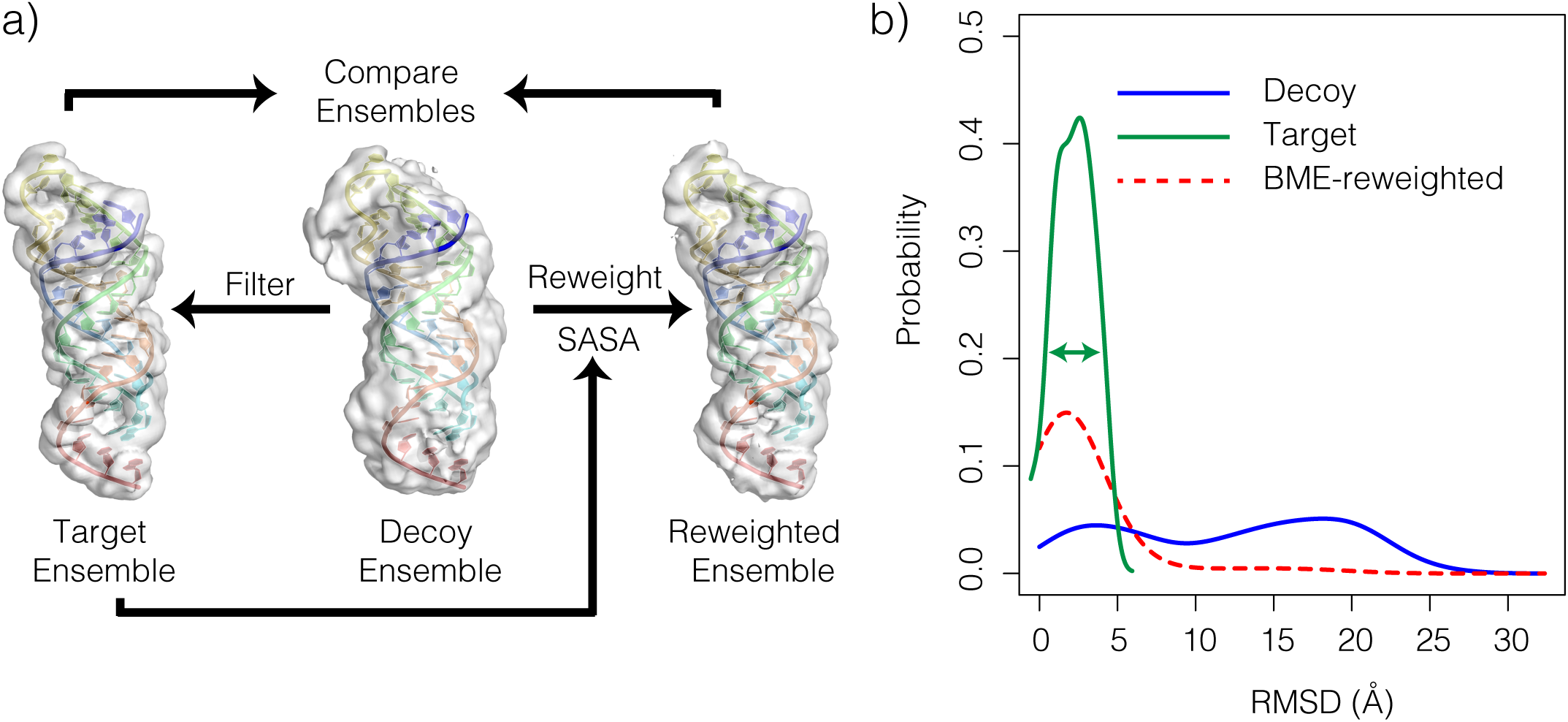
(a) Illustration of the workflow used to test the ability of SASA data to infer atomic ensembles. A target ensemble is constructed from a decoy ensemble by filtering structures based on RMSD from a reference structure. SASA data calculated from the target ensemble is used to reweight the decoy ensemble, and then the reweighted ensemble is compared to the target ensemble. (b) Distribution plots highlight the expected effect of reweighting a decoy ensemble using BME. Data from the data target ensemble is used to reweight the conformers in a decoy ensemble such that some distribution computed from the reweighted ensemble (*red*) more closely resembles the target distribution (*green*) than does decoy distribution (*blue*).

## Adding noise to target SASA data

As a more stringent test of the power of C8-SASA in ensemble reweighting, we carried out tests with target SASA data that included random noise. By adding noise to target SASA, we attempted to mimic SASA data that would be derived from experimental observables which contain systematic or statistical error at various levels and exhibit strong but imperfect correlation with SASA. We sampled random white noise from an exponential distribution with mean ranging from 0.3 to 4.8 Å^2^ with a random sign, and added it on to target SASA value for each atom. Note that the mean of error of 1.27 Å^2^ corresponded to the SASA data derived from LASER experiments for SAM riboswitch.

To mimic LASER-derived C8-SASA, we added random white noise to computed C8-SASA with mean of 1.27 Å^2^. The value of 1.27 Å^2^ was estimated from the error of LASER derived C8-SASA. We made a linear fit on computed SASA to LASER reactivity, and the absolute value of error of the fit was best estimated by an exponential distribution with scale parameter *β* = 1.27 (Figure S2). The resulting linear fit had a correlation score of 0.535 (and the mean absolute error was 2.69 Å^2^).

**Figure S2:**
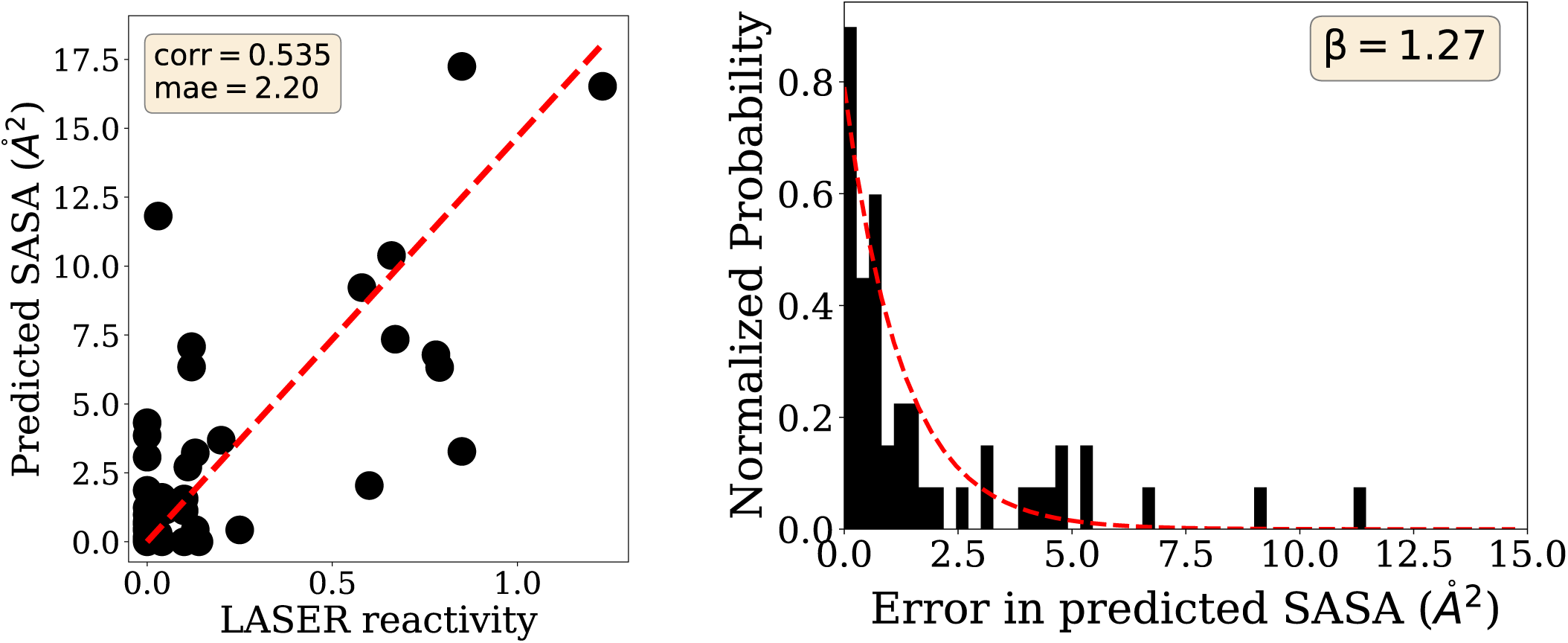
Linear Fit of C8-SASA to LASER reactivity. Left: the linear fit of C8-SASA to LASER reactivity. Right: exponential distribution of absolute error of the linear fit

## Difference atomic density maps for benchmark RNAs

To visually inspect individual ensembles, we calculated the difference between the atomic density maps (diffmaps) of the target (*T*) and the decoy (*D*) ensembles and compared them to the diff maps of the target (*T*) and the reweighted (*R*) ensembles. Shown in Figure S3 are comparisons between difference atomic maps and their corresponding cross correlations (*CC*) for all benchmarking RNAs between (left) the target and decoy ensembles (*CC*_*DT*_), (middle) the target and reweighted ensembles without (middle, *CC*_*RT*_) and (right) with noise (*CC*_*RT* (*noisy*)_). The diff maps are rendered in PyMOL as *fofc* volume maps, in which green and red color indicate -/+ 3 sigma contours. A higher *CC* value suggests the two maps in comparison closer resemble each other. Converly, more apparent green and red color in diff maps indicates the difference between the two maps are more prominent. Shown are target ensembles with RMSD width 2.2Å and in case of reweighting with noise, the noisy level is set to *β* = 1.27 which matches the error level in LASER-derived SASA for SAM-riboswitch.

Analysis of the difference maps indicated that in contrast to decoy ensembles (for example, Figure S3(1)(left)), the ensembles reweighted with noise-free SASA generally exhibited small residual densities relative to the target densities (for example, Figure S3(1)(middle)), confirming that conformational distribution in these ensembles closely mirrored the target. Ensembles that were reweighted with SASA data that contained simulated noise exhibited larger residual densities relative to target ensembles than their corresponding noise-free counterparts, but smaller residual densities than decoy ensembles (for example, Figure S3(1)(right)). These results provide a visual confirmation that, indeed, the C8-SASA can be used to reweight the decoy ensembles toward the target from which the SASA was calculated. We note, however, that when noise is added to the target C8-SASA data, the reweighted ensembles do not perfectly recapitulate the target ensemble.

**Table S1:** Benchmark Dataset

| PDB ID | Chain Length | Purine Length | Coverage |
| --- | --- | --- | --- |
| 1KKA | 17 | 10 | 0.588 |
| 1L1W | 29 | 17 | 0.586 |
| 1LDZ | 30 | 19 | 0.633 |
| 1NC0 | 24 | 13 | 0.542 |
| 1OW9 | 23 | 16 | 0.696 |
| 1PJY | 22 | 12 | 0.545 |
| 1R7W | 34 | 16 | 0.471 |
| 1SCL | 29 | 17 | 0.586 |
| 1UUU | 19 | 8 | 0.421 |
| 1XHP | 32 | 16 | 0.500 |
| 1YSV | 27 | 14 | 0.519 |
| 1ZC5 | 41 | 23 | 0.561 |
| 2FDT | 36 | 18 | 0.500 |
| 2JWV | 29 | 16 | 0.552 |
| 2K66 | 22 | 13 | 0.591 |
| 2KOC | 14 | 6 | 0.429 |
| 2L3E | 35 | 15 | 0.429 |
| 2LBJ | 17 | 8 | 0.471 |
| 2LBL | 17 | 7 | 0.412 |
| 2LDT | 31 | 18 | 0.581 |
| 2LHP | 37 | 19 | 0.514 |
| 2LK3 | 24 | 14 | 0.583 |
| 2LP9 | 16 | 9 | 0.562 |
| 2LPA | 15 | 8 | 0.533 |
| 2LQZ | 27 | 12 | 0.444 |
| 2LU0 | 49 | 28 | 0.571 |
| 2LUB | 37 | 19 | 0.514 |
| 2LUN | 28 | 14 | 0.500 |
| 2LV0 | 24 | 14 | 0.583 |
| 2M12 | 23 | 14 | 0.609 |
| 2M21 | 21 | 9 | 0.429 |
| 2M22 | 23 | 13 | 0.565 |
| 2M24 | 29 | 15 | 0.517 |
| 2M4W | 17 | 11 | 0.647 |
| 2M5U | 22 | 11 | 0.500 |
| 2M8K | 48 | 18 | 0.375 |
| 2MEQ | 19 | 10 | 0.526 |
| 2MFD | 19 | 9 | 0.474 |
| 2MHI | 53 | 24 | 0.453 |
| 2MNC | 29 | 14 | 0.483 |
| 2N2O | 23 | 11 | 0.478 |
| 2N2P | 23 | 11 | 0.478 |
| 2QH2 | 24 | 12 | 0.500 |
| 2QH4 | 18 | 10 | 0.556 |
| 2Y95 | 14 | 7 | 0.500 |

**Figure S3:**
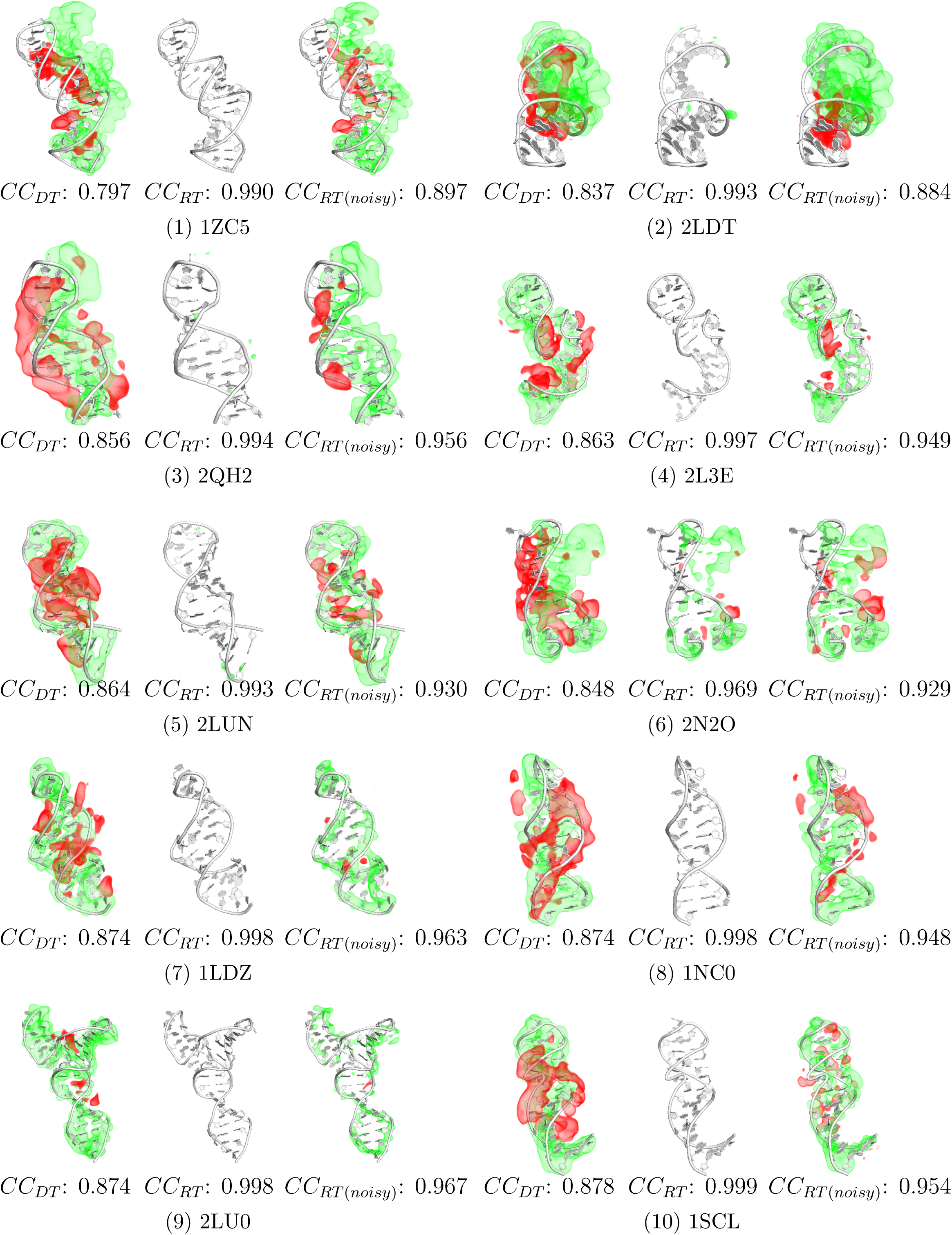
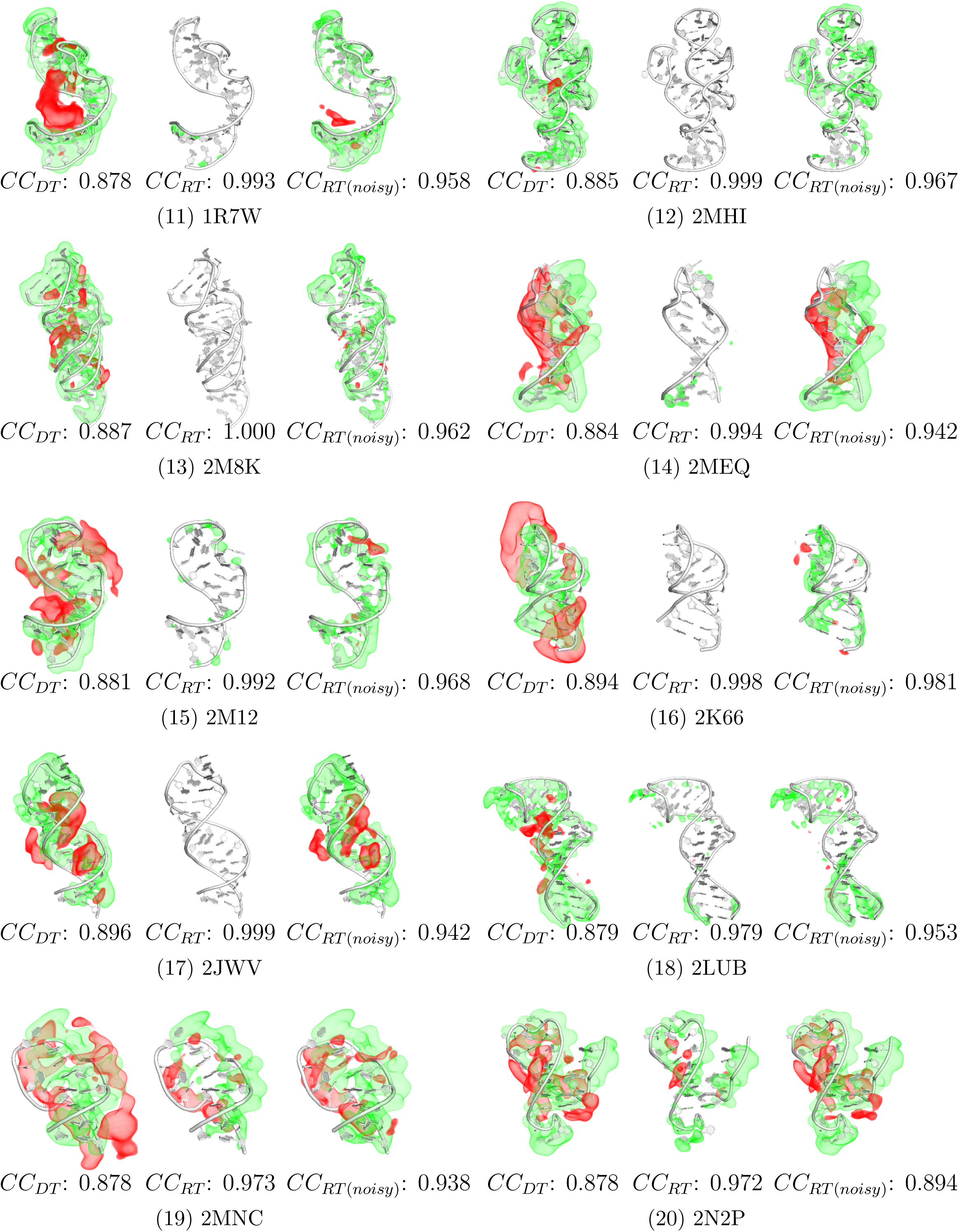
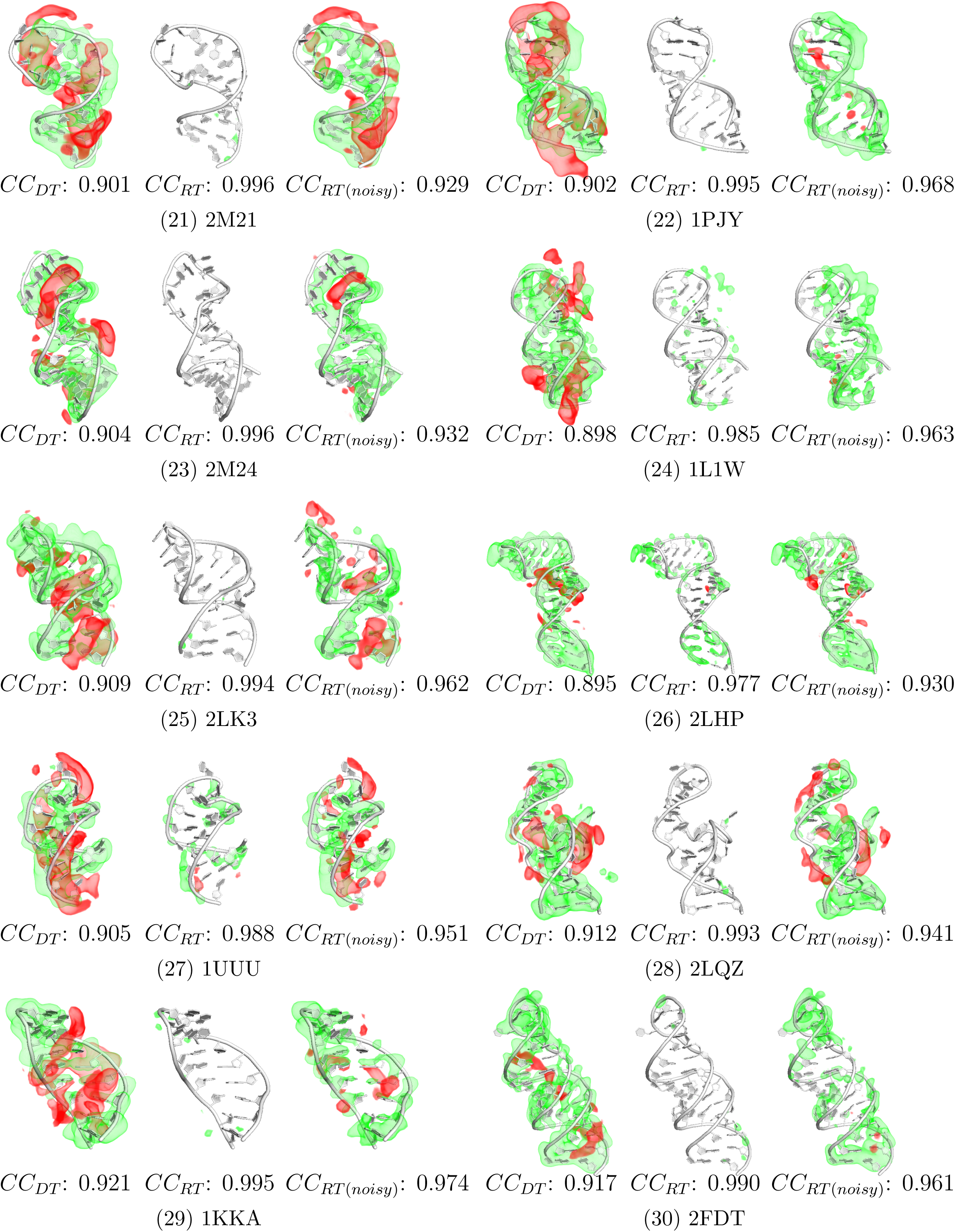
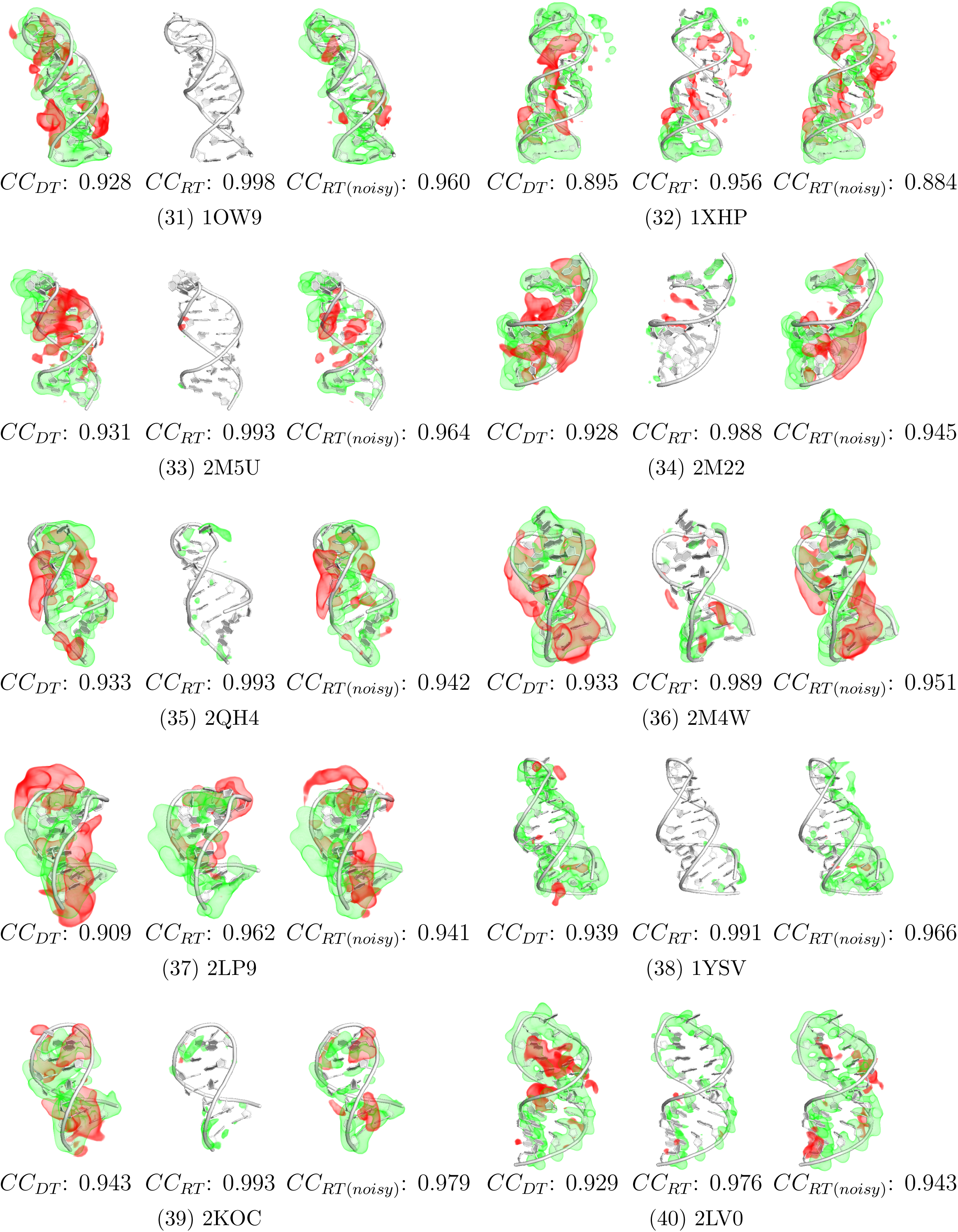
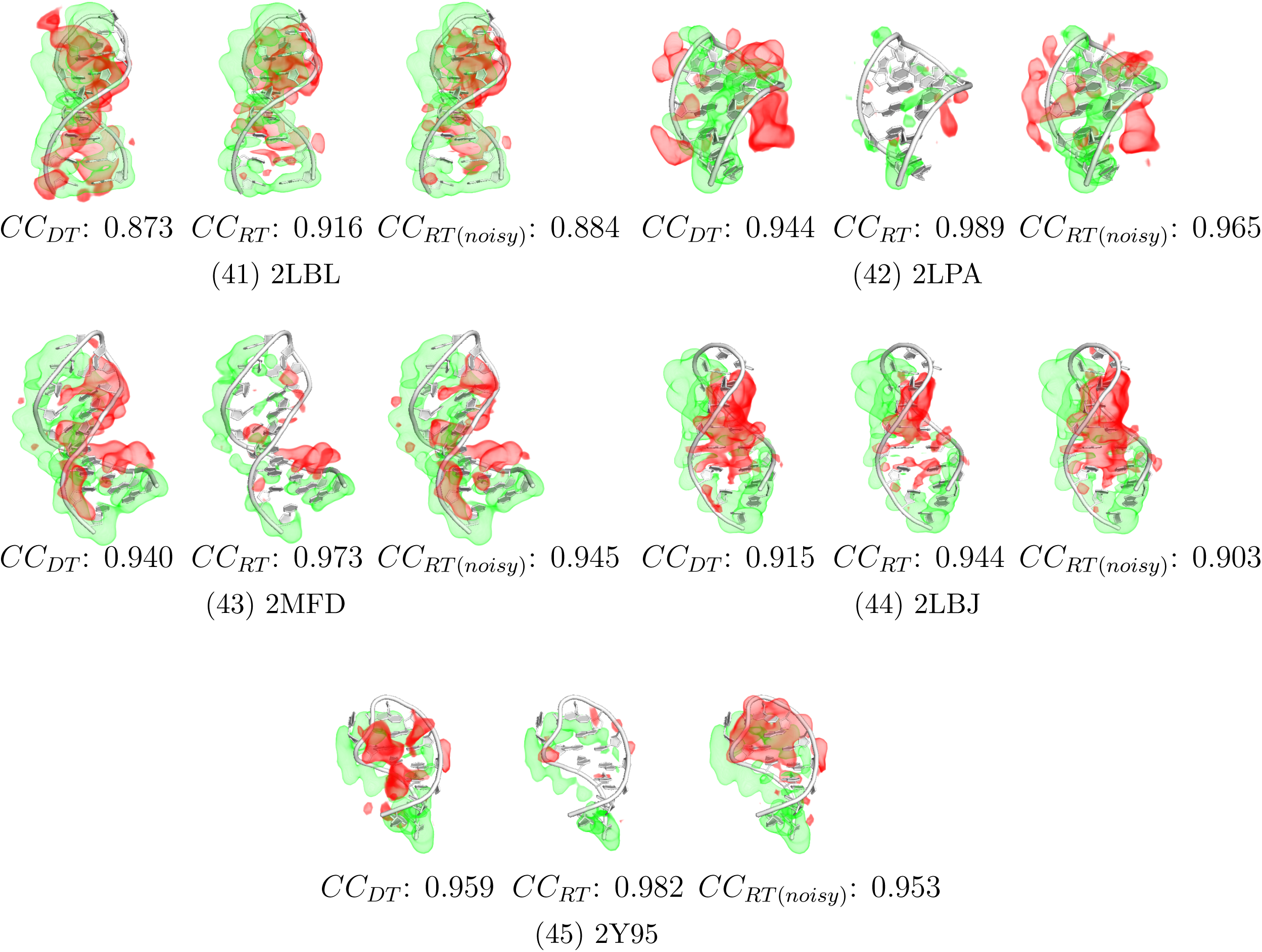
Difference atomic maps.

**Figure S4:**
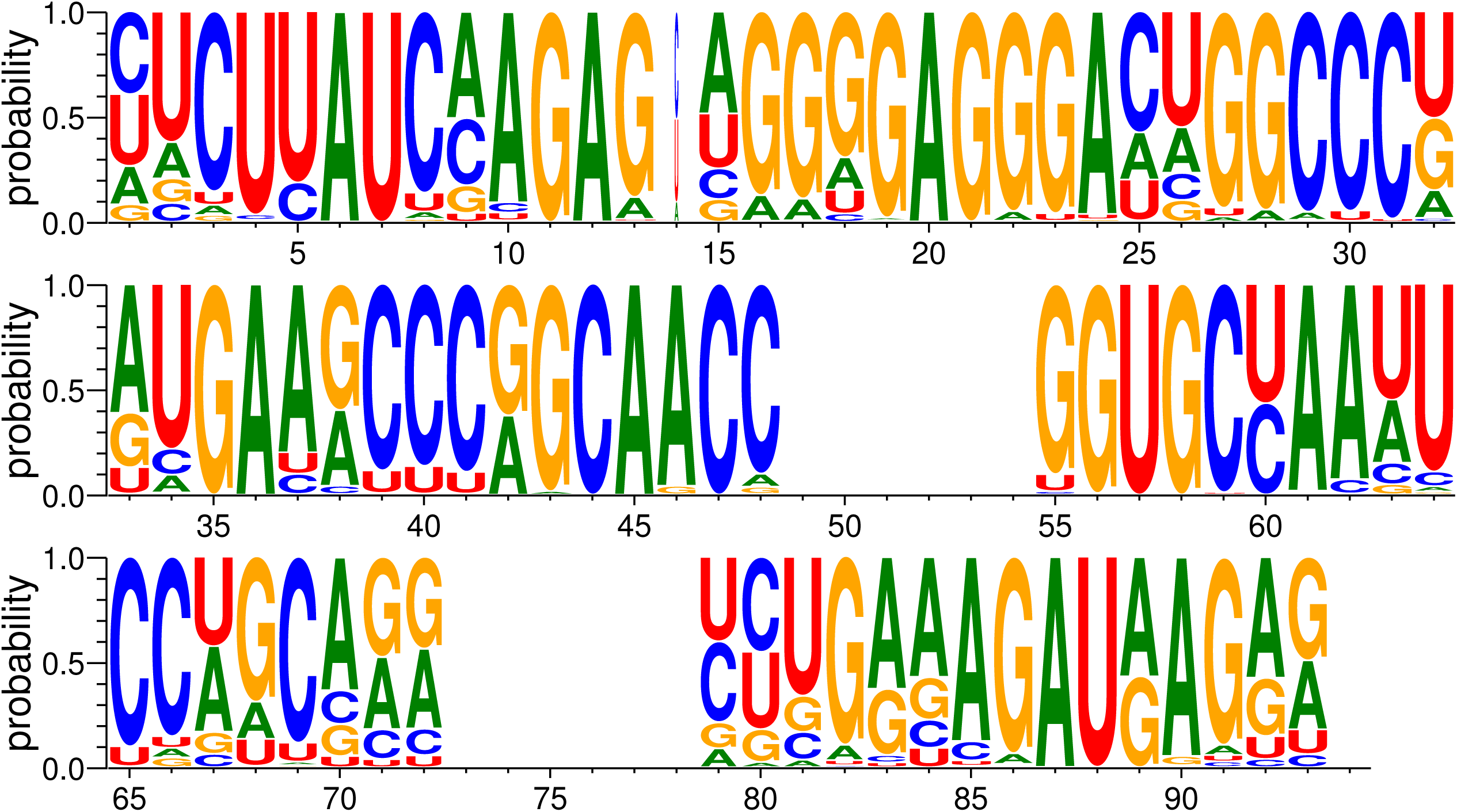
Conservation map for SAM-I riboswitch.

